# Cyanobacteria from marine oxygen deficient zones encode both form I and form II rubiscos

**DOI:** 10.1101/2024.09.06.611767

**Authors:** Alexander L. Jaffe, Kaitlin Harrison, Navami Jain, Leah J. Taylor-Kearney, Renée Z. Wang, Noam Prywes, Patrick M. Shih, Jodi Young, Gabrielle Rocap, Anne E. Dekas

## Abstract

Cyanobacteria are highly abundant in the marine photic zone and primary drivers of the conversion of inorganic carbon to biomass. To date, all studied Cyanobacterial lineages encode carbon fixation machinery hinged upon form I rubisco enzymes within a CO_2_-concentrating carboxysome. Here, we report that the AMZ IB lineage of *Prochlorococcus* from global oxygen deficient zones (ODZs) harbor both form I and form II rubisco enzymes, the latter of which are typically non-carboxysomal and possess biochemical properties tuned towards low oxygen environments. Our analyses reveal that these cyanobacterial form II enzymes are functional *in vitro* and were likely acquired via lateral gene transfer from proteobacteria. Global metagenomic read recruitment demonstrates that *Prochlorococcus* with form II rubisco are essentially restricted to ODZs in the Eastern Tropical Pacific, suggesting that acquisition may confer an advantage specifically under low-O_2_ conditions. Populations of AMZ IB *Prochlorococcus* express both forms of rubisco *in situ*, with the highest form II rubisco expression at depths where both oxygen and light are particularly low, possibly as a mechanism to increase the efficiency of photoautotrophy under energy limitation. Our findings expand the diversity of carbon fixation configurations in the microbial world and may have implications for the overall capacity of ODZs to sequester carbon.

## Introduction

The Calvin-Benson-Bassham (CBB) cycle is the dominant mechanism by which inorganic carbon is fixed biologically on Earth (1, 2). All members of the photoautotrophic phylum Cyanobacteria perform the CBB cycle using form I ribulose 1,5-bisphosphate carboxylase/oxygenase (rubisco) ensconced in a carboxysome, a bacterial microcompartment that serves as a CO_2_-concentrating mechanism. In the marine environment, the numerically abundant picocyanobacterial lineages *Prochlorococcus* and *Synechococcus* are thought to exclusively employ a form IA rubisco and α-carboxysomes, both of which may have been laterally transferred from proteobacteria (3). Among these groups, little variation within the genetic architecture of the carbon fixation machinery has been observed, despite the existence of numerous ecotypes differentiated by their spatial and metabolic niches (4). Here, we characterize the first known form II rubisco in Cyanobacteria, found within a highly abundant *Prochlorococcus* ecotype from global oxygen deficient zones (ODZs) (5, 6). Drawing upon a combination of metagenomics, metatranscriptomics, and metaproteomics, we also explore the ecological significance of this gene inventory and consider its potential ramifications for the marine carbon cycle.

## Results

While conducting a metagenomic survey of autotrophic organisms in the marine environment (7), we detected genomic bins from cyanobacteria encoding both form I and form II rubisco. Initial phylogenetic reconstructions indicate that these genomic bins fall within the AMZ lineages of *Prochlorococcus*, a set of marine ecotypes recently characterized from single-cell genomes (SAGs) from anoxic waters (5). To increase the small number of existing genomes for these lineages, we reconstructed 54 new medium-to-high quality metagenome-assembled genomes (MAGs) from public data. Analysis of genetic inventories of the total genome set indicated that only the AMZ IB subclade encoded both the form I and form II rubiscos in the same MAG/SAG (Fig. 1a, Supporting dataset 1). Although a fraction of AMZ IB possesses only the form I or form II, this is likely due to genome incompleteness.

**Figure 1.**
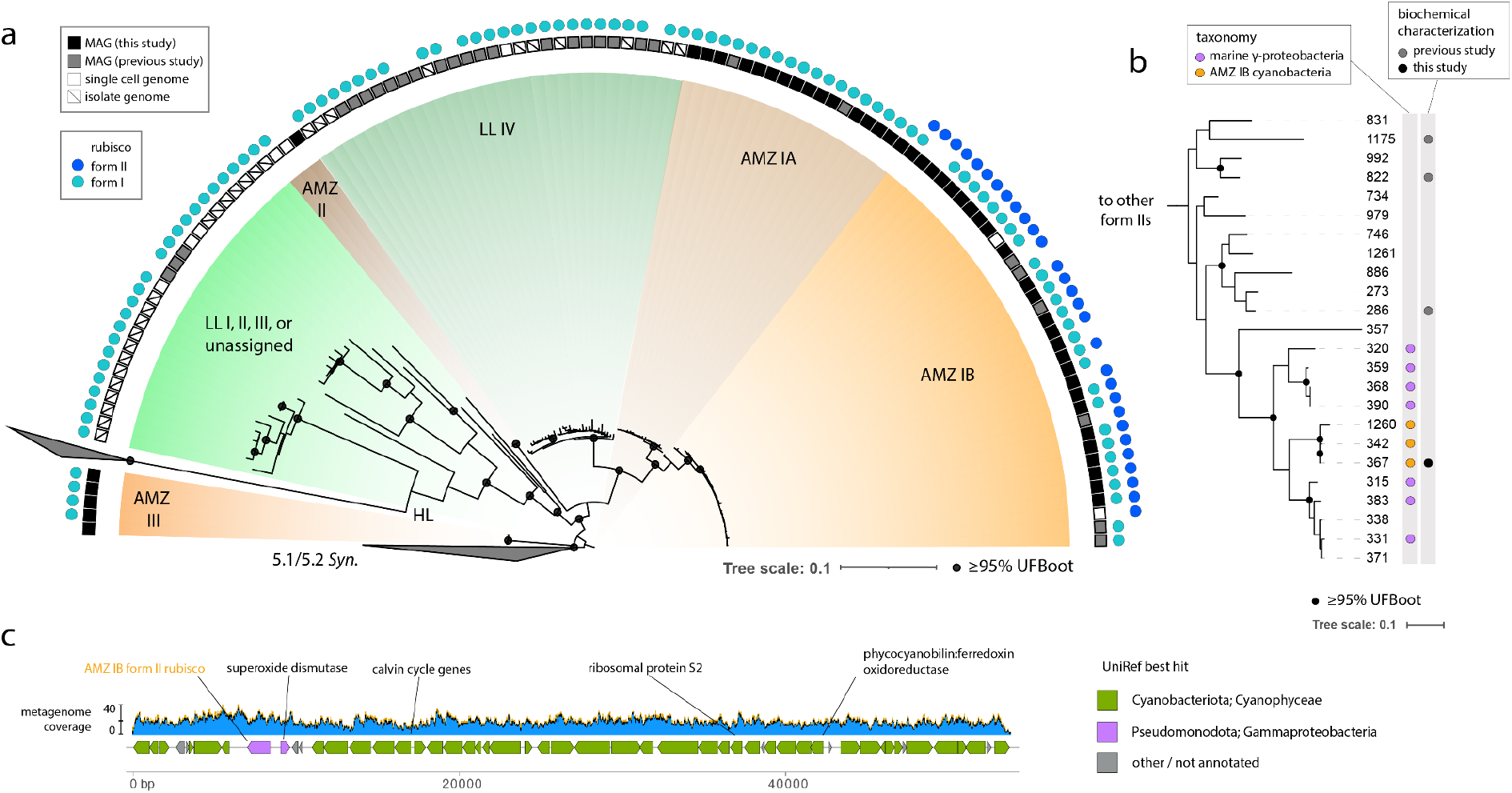
Genomic and phylogenetic characteristics of Cyanobacteria with multiple rubisco forms. **a)** Ribosomal protein tree depicting ODZ *Prochlorococcus* lineages and their rubisco inventories. Marine *Synechococcus* lineages (5.1-2) are used as outgroups. **N.B**. Redundant AMZ genomes are included for visualization purposes. **b)** Form II rubisco gene tree depicting representative AMZ IB sequences and their phylogenetic context. Tree tips are labeled with sequence numbers and relevant metadata (Supporting dataset 2). **c)** Genomic context of form II rubisco within a representative AMZ IB MAG. Blue regions represent the depth of metagenomic sequencing coverage along the contig.

We next clustered cyanobacterial form II rubiscos and placed select representative sequences into a protein tree. Representative sequences formed a small monophyletic clade within a larger group derived from the gammaproteobacteria, some of which were similarly isolated from seawater (Fig. 1b). We inferred that the cyanobacterial sequences were likely functional based on the proximity of other biochemically characterized proteins (Fig. 1b) and the presence of all conserved active site residues that coordinate substrate binding and catalysis (8, 9) (Supporting dataset 3). To confirm functionality, we heterologously expressed the most common *Prochlorococcus* form II sequence detected here (#367) and demonstrated its activity coupled to NADH oxidation (Extended Methods). The measured rate constant (*k*_*cat*_) was 1.2 ± 0.02 s^-1^, within the known range of measured form II rubiscos (10).

Finally, we examined the genomic context of cyanobacterial form II sequences, finding that they are encoded alongside a superoxide dismutase also of putative gammaproteobacterial origin (Fig. 1c, Supporting dataset 4), and were recovered on distinct scaffolds from form I sequences. No chaperone proteins or regulators were identified in the immediate context. Importantly, the overwhelming cyanobacterial affiliation of genes near the form II - most notably, the ribosomal protein S2 - indicate that the presence of form II rubisco is likely not the result of mis-binning; similarly, consistent read coverage indicates that mis-assembly in these regions is unlikely.

In addition to carbon dioxide, rubisco also reacts promiscuously with oxygen, leading to net carbon loss through a process termed photorespiration (11). Because form IIs generally possess a lower specificity for CO_2_ over O_2_ as a substrate (S_C/O_) and are not associated with carboxysomes, they are thought to be preferentially employed in low oxygen environments (12). Accordingly, we next examined the distribution of AMZ IB with form II rubiscos using a global metagenomic survey (Extended Methods). Our analysis detected AMZ IB within nearly 80 samples, essentially restricted to ODZs of the Eastern Pacific (Fig. 2a, Supporting dataset 5).

**Figure 2.**
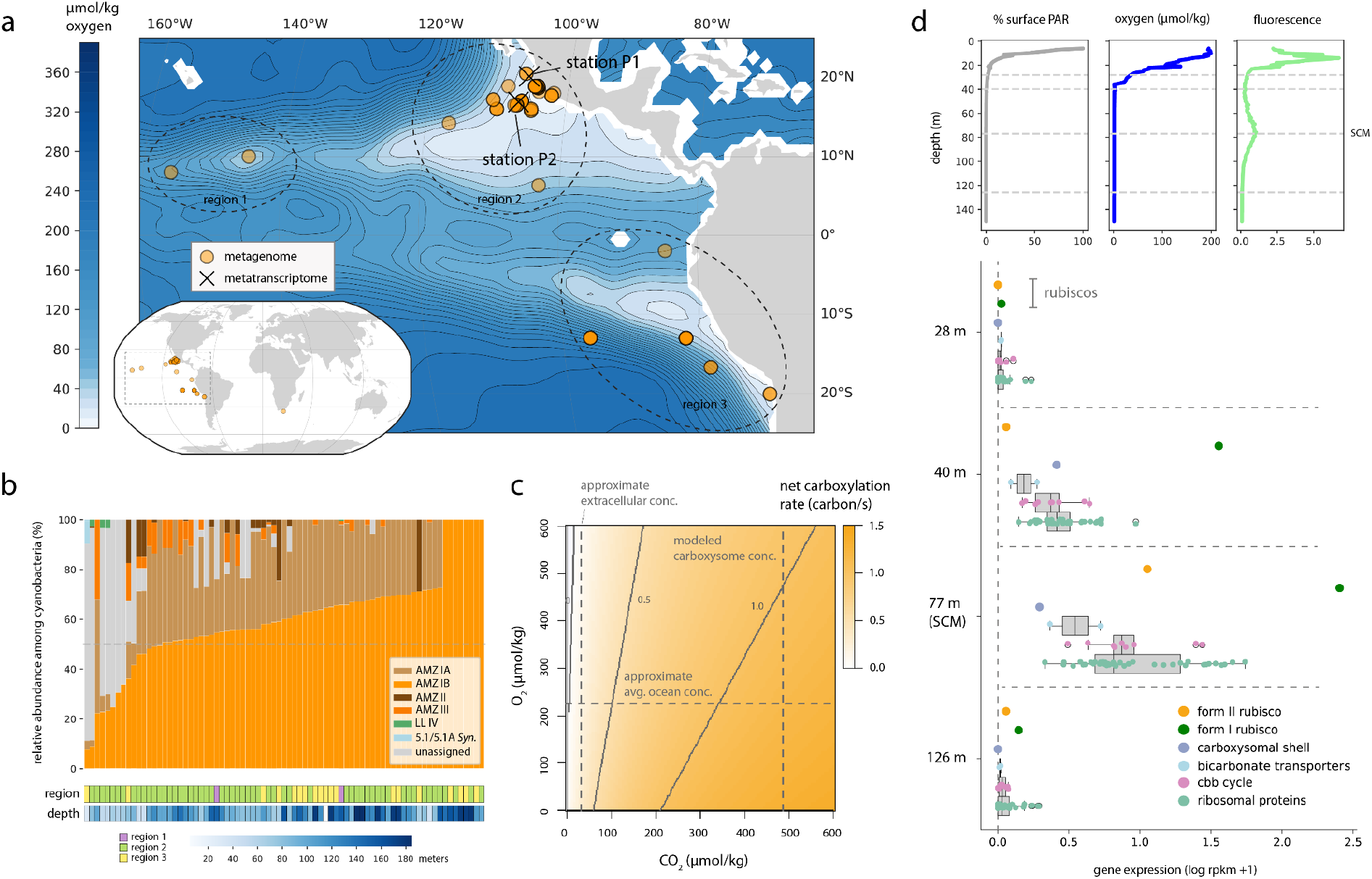
Distribution and expression patterns of AMZ IB *Prochlorococcus*. **a)** Geographic location of metagenomic and metatranscriptomic samples in which AMZ IB with form II rubisco were detected, overlaid with oxygen concentrations at 125 m (World Ocean Atlas 2018). **b)** Relative abundance (based on sequencing coverage) of cyanobacterial ecotypes in metagenomes (≤200 m) containing AMZ IB *Prochlorococcus*, with sample region and depth. **c)** Biochemical model estimating the net carboxylation rate of a representative form II rubisco (*T. denitrificans)* at 15°C as a function of intracellular carbon dioxide and oxygen. **d)** Environmental profiles and AMZ IB gene expression patterns with depth at station P1 in the Northeast Pacific. Each point in the lower panel represents the transcription level of a single gene. Abbreviations: PAR, photosynthetically active radiation. SCM, secondary chlorophyll max.

When detected, this lineage typically comprised the majority of the cyanobacterial community (Fig. 2b), consistent with previous results (5, 6). Supporting this distribution, biochemical modeling using the kinetic features (k_cat,C_, K_C_, K_o_, S_C/O_) of a representative form II rubisco (*T. denitrificans*) with a similar rate constant to that of AMZ IB suggests that net carboxylation occurs at typical ODZ conditions, assuming that intracellular concentrations of CO_2_ fall in between those of seawater and the carboxysome interior (Fig. 2c, Extended Methods).

Within oxygen-deficient zones (ODZs), oxygen concentrations, light, and nutrients vary significantly with water depth (6). To examine the impact of the gradients upon the utilization of different rubisco forms, we leveraged a depth-resolved metatranscriptomic transect from an offshore ODZ near Mexico (station P1, Fig. 2a) (13). Alignment of sequenced transcripts to a representative genome suggested that AMZ IB was only active in the lower portion of the ODZ euphotic zone (Fig. 2d). Expression of the AMZ IB form I rubisco was very high at both 40 m and 77 m, whereas the form II was essentially only expressed at 77 m (the anoxic secondary chlorophyll max, ∼0.01% surface PAR and ∼1.09 μmol/kg O_2_), where it attained approximately 4% of form I levels. Trends were similar at another site (P2) with a more gradual oxycline and deeper SCM, though relative expression of the form II was higher (∼16% of form I at the SCM) (Fig. 2a, Supporting dataset 6). Additionally, we detected 10 peptides identical to the predicted AMZ IB sequence in a metaproteome previously collected at P2 (6) (Supporting dataset 7). These peptides were only recovered at 100 meters, below the oxycline near the top of the SCM.

## Discussion

Here, we demonstrate that a highly abundant lineage of cyanobacteria from ODZs uncharacteristically encode (and express) both the form I and form II rubisco. While known for a number of marine proteobacteria, this dual genetic repertoire has not before been observed in Cyanobacteria, Earth’s most numerous bacterial primary producers. In proteobacteria, form II rubiscos are cytoplasmic (non-carboxysomal) and can be preferentially transcribed under low oxygen conditions that theoretically favor this enzyme form’s biochemical attributes (14, 15).

Thus, we propose that the acquisition of the form II rubisco by AMZ IB - likely from co-occurring gammaproteobacteria - represents an adaptation to low oxygen niches within ODZs. This gene, alongside others related to anaerobic stress tolerance and signaling (5), may in part permit AMZ IB to flourish in the measurably anoxic waters of the euphotic zone.

At this stage, the physiological impact of simultaneous rubisco expression in AMZ IB is unclear. One possibility is that tandem expression permits cells to fix additional CO_2_, including that leaked from the carboxysome associated with their form IA rubisco. We speculate that the energetic cost associated with carboxysome production may be particularly burdensome at depths where light (and thus energy) is limited, and thus that a non-carboxysomal form might be particularly advantageous there. Tandem expression of both forms could also be explained in part by a dependence of the form II rubisco upon form I-associated chaperones or regulators.

Together, our results demonstrate a novel metabolic configuration with the potential to impact carbon cycling in ODZs, which are significant and expanding portions of the world’s oceans. Additionally, our findings raise the intriguing possibilities that conserved carbon fixation genes may be more mobile and modular than previously recognized, and that harboring multiple forms of rubisco might be a viable strategy towards increasing carbon fixation efficiency.

## Materials and Methods

Materials and methods are available in the Supporting Information.

## Data, Materials, and Software Availability

All underlying data are available in the Supporting Information or, in the case of newly-resolved AMZ MAGs, through the NCBI under Bioproject accession PRJNA1136951. Custom code used for the analysis of all data is available at github.com/alexanderjaffe/odz-pros.

## Acknowledgements

We thank Christina Rathwell, Natalie Kellogg, Alex Roberts, Luis Valentin-Alvarado, and Matthew Olm for aid with data access and interpretation. We thank the Stanford Research Computing Center and the Stanford Geomicrobiology Shared Laboratories Core Facility (RRID:SCR_025000). Funding was provided by the Stanford Science Fellows program, the NSF Postdoctoral Fellowship in Ocean Sciences (A.L.J.), NSF award OCE-2143035 (A.E.D.), NSF award OCE-2022911 (G.R.), and the Simons Foundation Award 561645 (J.Y.).

## Notes

### Competing Interest Statement

The authors have declared no competing interest.

