## Supplementary Information for "Cyanobacteria from marine oxygen deficient zones encode both form I and form II rubiscos"

**This PDF file includes:**

Extended methods
Extended methods references

**Other supporting materials for this manuscript include the following:**

Supporting datasets 1-7

### Extended Methods

**Initial genomic database construction.** Cyanobacterial genomes putatively encoding the form II rubisco were taken from a recent study reporting genome information and gene inventories for a large number of rubisco-encoding organisms (1). Preliminary ANI-based comparisons indicated that these genomes were related to a subset of lineages previously described from a set of single-amplified genomes (SAGs) from anoxic marine zones in the Eastern Tropical Pacific (2). The total set of SAGs from this study, as well as reference sequences, were downloaded from NCBI or the Integrated Microbial Genomes (IMG) portal and combined with initial genomes.

Observing a paucity of high-quality genomes specifically associated with the AMZ IA, IB, II, III clades, we attempted to recover additional sequences through a targeted metagenomics effort. First, the highest quality representative sequences for each of the four AMZ lineages were determined by CheckM (3) and uploaded to the Branchwater web interface (https://branchwater.jgi.doe.gov/). Resulting files described the incidence of cyanobacterial lineages of interest across the millions of global metagenomes indexed by Branchwater and were subsequently filtered to 0.97 cANI, which frequently indicates a 'species' level match. Bioproject accessions for these samples were recorded, and those Bioprojects with four or more positive samples (4–9) were retained for further analysis.

To recover new metagenome-assembled genomes (MAGs) for species of interest, we subjected the samples identified above to a typical metagenomics workflow implemented in Snakemake (10) (available at github.com/alexanderjaffe/odz-pros). Briefly, raw metagenomic reads were downloaded from the Sequence Read Archive using parallel-fastq-dump (github.com/rvalieris/parallel-fastq-dump), screened for mispaired reads using repair.sh from the BBMap tool suite (sourceforge.net/projects/bbmap/), and further quality-filtered using bbduk.sh from the same tool suite. Trimmed reads were assembled using MEGAHIT v. 1.2.9 (11), and resulting contigs were indexed for read alignment using bowtie2 (12). Within each Bioproject, mapping proceeded in an 'all-by-all' fashion, whereby contigs from each sample were aligned to trimmed metagenomic reads from every other sample within that same Bioproject. This alignment matrix was passed to metabat2 (13) for differential coverage binning.

Resulting bins that affiliated with the Cyanobacteria were identified using GTDB-Tk (14) and compared to the initial genome set using dRep (15). Newly-resolved bins that clustered with any AMZ lineage at 95% average nucleotide identity (ANI) and met lax quality criteria (≥40% completeness and ≤25% redundancy) were retained. Preliminary lineage assignments were also made for new genomes based on ANI.

**Re-assembly and curation of metagenome-assembled genomes.** Secondary assembly was repeated for each newly-resolved bin using the subset of reads that mapped to it in the original metagenomic assembly. This process was hypothesized to reduce fragmentation and increase the length of resulting contigs. Mapped reads were extracted from BAM files using samtools, converted to fastq files with seqtk, and re-assembled with metaspades (v3.15.5 and default parameters). Quality of re-assembled genomes was measured by a) the number of contigs with length over 1500 bp, b) N50, and c) total length of the genome assembled. New versions were chosen if and only if scaffold count decreased, N50 increased, and total length decreased by no more than 10% of the original. Otherwise, the original was retained.

Bins were further refined using an approach that identifies putatively contaminating scaffolds based on aberrant protein content (16). Briefly, proteins were predicted for all AMZ lineages (IA, IB, II, and III) using Prodigal (17) and clustered into protein families on a per-lineage basis using MMseqs2 (*--cov-mode 0*)

(18). A protein family was considered ‘rare’ if it was detected in less than 10% of genomes from a given lineage. Contigs composed of 50% or more rare protein families were removed.

**Phylogenomics of *Prochlorococcus* ecotypes.** The total set of genomes (including SAGs, newly-resolved MAGs, and other references) were secondarily filtered to more stringent quality threshold (≥50% completeness and ≤10% redundancy) and clustered again at 99% ANI using dRep. For non-AMZ genomes, a representative was chosen on the basis of CheckM completeness. For visualization purposes, all AMZ genomes were utilized regardless of redundancy. Proteins were predicted using prodigal (17) and annotated using kofamscan (19); hits to HMM models with e-values ≤1e-5 were retained. These annotations were queried for the 16 syntenic ribosomal proteins frequently used for phylogenetic reconstruction from fragmented genome assemblies (20); in cases where multiple instances of a single protein were recovered, we prioritized the one on the most common scaffold. We further required that genomes have at least four of the expected 16 ribosomal proteins to be included in the tree.

Each phylogenetic marker was aligned individually using mafft (21) and subsequently trimmed using trimal (*-gt 0.1) (22)*. Individual alignments were concatenated on a per-genome basis and passed to IQ-Tree2 (*-m TEST -st AA -bb 1000)* (23) for maximum-likelihood tree inference with 1000 ultrafast bootstraps. Marine *Synechococcus* lineages 5.1 and 5.2 were used as outgroups, based on a previously published species tree topology for AMZ clades (2).

**Classification and molecular phylogenetics of rubisco gene inventories.** Protein predictions for each genome meeting quality thresholds were searched for rubisco using published form-specific hidden markov models (HMMs) (1). HMM results were read into Python using the SearchIO package and filtered such that query sequences covered at least 50% of the HMM model. If a query attained above-threshold hits to multiple models, the model with the highest score was chosen. Rubisco gene inventories were visualized in tandem with the species tree resolved above using iTOL (24).

To establish a reference set of form II rubisco, protein sequences were gathered from three previous publications (25–27). To account for redundancy within the combined set, sequences were clustered using usearch at 100% identity (*-id 1 -sort length*) (28). Within each cluster, we curated sequence metadata concerning species of origin, oligomeric state (if measured), and catalytic rate (if measured) (Supporting dataset 2). Representative proteins from each cluster were aligned and subjected to phylogenetic reconstruction as above if they attained ≥50% of alignment length.

**Measurement of form II kinetics.** Synthesis, purification, and *in vitro* kinetic assays followed those in Davidi et al. 2020 (26). Briefly, the amino acid sequence for most common AMZ IB form II sequence (#367) was synthesized into the pET28-14xHis-bdSumo expression vector (26)(Twist Biosciences) and transformed into BL21(DE3) Star *E. coli* competent cells (QB3 Macrolab) *via* heat shock. Starter cultures were then grown to high density in LB with Kanamycin (Apex) (37°C/200 RPM, overnight). An aliquot of this culture was mini prepped and sent for Primordium sequencing to confirm the ORF sequence. 50 mL of LB was inoculated with the starter culture and grown to high optical densities (OD600~0.85), upon which rubisco expression was induced with 1 mM IPTG (Apex) for ~16 hours (16 °C/200 rpm).

Cells were then pelleted (5,000 rpm, 20 min, 4°C), subject to a freeze-thaw cycle (-80 C), then thawed and lysed with BugBuster (Millipore) in the presence of DNAse. Cell lysate was clarified by centrifugation (14,000g at 4 °C for 20 min). The soluble fraction was applied to Pureproteome Nickel Magnetic beads (Millipore) for batch binding. Beads were washed twice with increasing concentrations of imidazole (25 and 50 mM imidazole; 20 mM HEPES; pH 8; 100 mM NaCl; 20 mM $MgCl_2$) followed by low imidazole concentration wash (5 mM) to remove excess imidazole. The beads were then resuspended in the low imidazole concentration buffer and rubisco was cleaved *via* application of SUMOlase enzyme for ~16 hours with gentle rocking at 4 °C. Cleaved protein was separated from the beads *via* centrifugation; purity was confirmed via SDS-PAGE and yield *via* Nanodrop.

The carboxylation rate of AMZ IB form II rubisco was measured using a previously published spectroscopic assay that couples activity to NADH oxidation (26, 29, 30). Here, assay components were mixed in a 96 well flat-bottom transparent plate (Corning, Costar) at pH 8, 25 °C, 4% $CO_2$ and 0.5% $O_2$ with orbital shaking at 1440 rpm. Rubisco was activated *via* incubation with $CO_2$ for ~20 minutes at the working conditions and the reaction was initiated through the addition of RuBP. Active site concentration was estimated using the known rubisco inhibitor, CABP, at n=2 using 4 CABP concentrations. Measured rubisco rates were plotted against CABP concentration, with the x-axis intercept proportional to rubisco active site concentration. $V_{max}$ (*n*=3) was divided by this x-intercept ([E]) to generate $k_{cat}$. Rubisco catalysis rates and CABP inhibition slopes were calculated using a custom script (10.5281/zenodo.7757660). A previously measured 'fast' rubisco (4_10) was measured in parallel as a positive control.

**Modeling of rubisco kinetics.** Carbon fixation rate modeling based on Harrison et al. *submitted (31)*. Briefly, the net carboxylation rate is modeled as:

$$R_C - 0.5 * R_O$$

where $R_C$ is the gross carboxylation rate of Rubisco:

$$R_C = k_{cat,C} * \frac{[CO_2]}{[CO_2] + K_C + K_C * \frac{[O_2]}{K_O}}$$

and $R_O$ is the gross oxygenation rate:

$$R_O = \frac{R_C * [O_2]}{S_{C/O} * [CO_2]}$$

$k_{cat,C}$ is the carboxylation rate of each active site ($CO_2$/s), $S_{C/O}$ is the specificity for $CO_2$ versus $O_2$, $K_C$ and $K_O$ are the half saturation constants for $CO_2$ and $O_2$, respectively, and $[CO_2]$ and $[O_2]$ are the concentrations of $CO_2$ and $O_2$, respectively. Temperature scaling is done by creating Arrhenius equations for each kinetic parameter ($k_{cat,C}$, $K_C$, $K_O$, and $k_{cat,O}$):

$$x(T) = e^{c - \frac{\Delta H}{RT}}$$

where R is the universal gas constant (0.00831 kJ/mol/K), ΔH is approximated from reported values in Galmés et al. 2016 (32) and Galmés et al. 2015 (33), and T is temperature in K. c is calculated by setting the equation equal to the reported value for each kinetic parameter.

Temperature for the comparison was set at 15°C based on *in situ* CTD data from stations P1 and P2 (see below). Form II kinetic values were approximated using known values from *Thiobacillus denitrificans* (34) since, of the form II enzymes for which all kinetic values are available, it has the $k_{cat}$ closest to the value measured in this study. Values used for this analysis are found in the table below.

| Parameter | | Form II rubisco (Approximation) | | |
|---|---|---|---|---|
| | | Value | Source | Ref |
| $k_{cat,C}$ | Value @25°C | 2.3 | *T. denitrificans* | (34) |
| | ΔH | 52.9 | Average of available values[1] | (32) |
| $K_C$ | Value @25°C | 256 | *T. denitrificans* | (34) |
| | ΔH | 40.8 | Average of available values[1] | (32) |
| $K_O$ | Value @25°C | 619 | *T. denitrificans* | (34) |
| | ΔH | 26.7 | Average of all in-vivo $K_O$ ΔH[2] | (32) |
| $S_{C/O}$ | Value @25°C | 13.3 | *T. denitrificans* | (34) |
| | ΔH | -18.8 | Value from *R. rubrum* form II | (32) |

[1]Excluding Rhodophyta, as Galmés et al. 2016 (32) notes how different it is from the others.
[2]Excluding *A. thaliana* because its $r^2$ value was much lower than the others.

Environmental $CO_2$ concentrations at station P1 were estimated using dissolved inorganic carbon (DIC) values from GO-SHIP Section P18 (NCEI Accession 0171546). A value for the $CO_2$ concentration inside the *Prochlorococcus* carboxysome was taken from Hopkinson et al. 2014 (35). Seawater $O_2$ concentrations were estimated from an average of World Ocean Atlas 2018 data points (125 m depth).

**Genomic context of form II rubiscos.** Proteins neighboring the cyanobacterial form II were subjected to a series of annotation steps to illuminate their functional and taxonomic associations. First, proteins were annotated using KEGG via kofamscan and Pfam (36) (e-value cutoff 1e-20). Next, proteins were subjected to a previously described BLAST-based workflow that compares sequences to UniRef100 (37) and assigns likely taxonomic affiliation using their best hit (1, 38). Context was visualized using gggenes.

**Ecological distribution and abundance of cyanobacterial ecotypes.** Representative genomes for all AMZ lineages, as well as any other co-occurring Cyanobacteria lineages, were selected from each 95% ANI (species group) cluster based on genome completeness and combined into an index for read mapping using bowtie2. We next aligned metagenomic reads from a subset of global marine metagenomes to this index using a previously published Snakemake workflow (1). This workflow was amended with one additional step to account for mispaired reads, and similar parameters for mapping specificity were employed (minimum 95% read identity, minimum mapq score of 20). Samples for mapping were selected if already shown to contain AMZ IB in the previous study (1) or were recovered by the Branchwater analysis described above (~90 samples total). As before, genomes were considered present if they attained a minimum coverage breadth of 50%, and at least half of the expected breadth as determined by inStrain (39). We also required that the AMZ IB form II rubisco gene itself attain at least 50% coverage breadth to be considered a true detection, which removed several samples. Cyanobacterial community composition was determined by summing all mean coverage values across representative genomes and determining the fraction of total coverage accounted for by each lineage.

To visualize the spatial distribution of the AMZ IB ecotype in the global ocean, geographic coordinates were obtained for each sample through a hybrid automated/manual approach. These coordinates were plotted onto a Robinson projection of a world map using cartopy (https://scitools.org.uk/cartopy/docs/latest/). Mean oxygen data from the 2018 World Ocean Atlas (40) were plotted as contours; oxygen concentrations at 125 meters were visualized.

**Gene expression patterns of AMZ IB in a coastal ODZ transect.** Gene expression analysis followed the workflow presented in (1), employing the non-redundant set of cyanobacterial genomes described above and a set of 16 previously published metatranscriptomes from a low-oxygen region in the Eastern Tropical North Pacific Ocean (41). Transcript abundances in reads per kilobase million (RPKM) were computed gene-wise for the representative AMZ IB genome (SRR11923207.100.13) and reconciled with functional annotations derived from kofamscan and Pfam (36). A subset of morning  samples from the onshore station P1 (20°9′0″N, 106°0′0″W) were selected for comparison. CTD data were obtained for the cast 59 – representing the P1 SCM sample – from Rolling Deck 2 Repository (https://www.rvdata.us/search/cruise/RR1805), averaged within 1 meter depth bins, and plotted.

**Supporting dataset 1.** Characteristics of genomes utilized in the study.

**Supporting dataset 2.** Sequence information and metadata for form II rubiscos utilized in the study.

**Supporting dataset 3.** Conservation of active site residues in AMZ IB form II rubisco. Column numbering follows *R. rubrum* protein sequence.

**Supporting dataset 4.** Genomic context for AMZ IB form II rubisco.

**Supporting dataset 5.** Characteristics of metagenomic samples used for relative abundance analysis.

**Supporting dataset 6.** Gene expression data for AMZ IB at stations P1-2.

**Supporting dataset 7.** Alignment of metaproteomic peptide fragments to AMZ IB form II rubisco.

### Extended methods references
